# Let’s talk about cardiac T1 mapping

**DOI:** 10.1101/343079

**Authors:** Tarik Hafyane, Agah Karakuzu, Catherine Duquette, François-Pierre Mongeon, Julien Cohen-Adad, Michael Jerosch-Herold, Matthias G. Friedrich, Nikola Stikov

## Abstract

**Background:** Recent reports have shown that T1 mapping sequences agree in phantoms, but exhibit significant differences in vivo. To characterize these differences in the heart, one needs to consider the effects of magnetization transfer (MT) and the T2 relaxation time in the most commonly used cardiac T1 mapping sequences (MOLLI, ShMOLLI and SASHA).

**Methods:** Six explanted pig hearts were scanned weekly over a period of six weeks on a 3T system with the MOLLI, ShMOLLI, SASHA sequences and an inversion recovery sequence as reference. The T1 bias was computed as the difference between MOLLI, ShMOLLI, SASHA and the reference T1 values. We applied robust correlation statistics to assess the relationships between T1, T2 and MT. All data are publicly available at: http://neuropoly.pub/pigHeartsData.

**Results:** A systematic T1 bias was present for all sequences, with MOLLI and ShMOLLI underestimating T1 and SASHA slightly overestimating T1 compared to the reference. The correlation of T1 bias with T2 was weak and insignificant. However, MT showed significant associations with T1 bias for all sequences. Our analysis is also available at: http://neuropoly.pub/pigHeartsInteractive.

**Conclusion:** We investigated cardiac T1 mapping sequences in a setting that allowed us to explore their accuracy and their dependence on T2 and MT effects. The T2 effects were not significant, and could not explain the T1 bias of MOLLI, ShMOLLI, SASHA with respect to the reference. On the other hand, the T1 biases exhibited a strong correlation with MT. We conclude that inaccuracies in cardiac T1 mapping are primarily due to magnetization transfer.

## BACKGROUND

Cardiac T1 mapping has the potential to play an important role in the diagnosis of heart disease ^1^. Abnormal native T1 values are indicators of infarction and inflammation ^2^, protein deposition ^3^, lipids ^4^, and iron accumulation ^5^. Abnormal post contrast T1 values are also consistent with extracellular volume expansion ^6^. T1 mapping in the heart is primarily performed via inversion recovery ^7^ or saturation recovery sequences ^8^, or a combination of the two ^9^. The modified Look-Locker sequence (MOLLI) ^10^ and its shortened version ShMOLLI ^11^ use magnetization inversion, and then sample it multiple times during its return to equilibrium. The saturation recovery sequence SASHA ^12^ uses saturation pulses (90 degrees) to minimize the longitudinal magnetization, and then samples it once during its return to equilibrium after saturation. All methods provide a T1 map from the same cardiac phase within a single breath hold.

T1 values obtained in normal volunteers show a significant difference between inversion recovery (MOLLI/ShMOLLI) and saturation recovery (SASHA) sequences ^13,14^. Bloch simulations to characterize the difference between the two methods point to the T2 parameter as a major bias that affects the MOLLI method and its variant ShMOLLI, whereas the SASHA sequence was minimally affected by T2 ^15^. Recent published Bloch simulations ^16,17^ found that in addition to T2, the magnetization transfer (MT) effect also introduces bias in MOLLI and its variant ShMOLLI, but not in SASHA. On the flipside, SASHA tends to produce noisier maps, seemingly trading off precision for accuracy ^14^.

The Society for Cardiovascular Magnetic Resonance (SCMR) and the CMR workshop group of the European Society of Cardiology have issued a consensus statement which mentioned Bloch simulations and phantoms as the first step in validating T1 mapping sequences, but recommended further research to study MT, T2 and T2* effects in biological tissues, as they may affect the T1 estimates differently compared to simulations and phantoms ^18^.

To study MT and T2 effects in a more realistic environment, we imaged ex vivo pig hearts (in addition to phantoms) and applied a robust statistical framework to characterize and interpret the differences we observe between phantoms and ex vivo measurements.

## METHODS

### Phantoms

The phantom used in this study was prepared by Captur and colleagues ^19^ and is composed of 3 3 array tubes with different agar concentrations (ranging from 0.2 to 3 %) and different T1/T2 values. T1 values in the phantom range from 250 to 3000 ms and T2 values from 50 to 150 ms. Only tubes with T2 values that correspond to the range of normal and abnormal myocardium T2 values were considered ^20,21^, hence a total of 6 tubes with a range of T2s from 50 to 65 ms were considered for the analysis.

### Ex vivo heart imaging

Six pig hearts were obtained from a local shop that supplies cardiology residents with specimens for anatomy lessons. The hearts were kept in ice during transportation, then immersed in saline for three hours before the first MRI scan. The hearts were imaged weekly (weeks 1, 2, 3) using a protocol approved by the institution’s research ethics board and described below. Subsequently the hearts were immersed in 10% neutral buffered formalin solution (Chaptec Inc., QC, Canada) and imaged for another three weeks (weeks 4, 5, 6) using the same MRI protocol.

### MRI protocol

T1, T2 and MTR measures were performed in a mid-ventricular slice of the ex-vivo hearts, and also in the agar phantoms using a 3T system (Magnetom Skyra, Siemens Healthcare, Erlangen, Germany) with an 18-channel phased-array cardiac coil. A simulated heart rate of 60 bpm was used. T1 measures were obtained in a single slice using the IR-TSE sequence as a reference, as well as the MOLLI, ShMOLLI and SASHA sequences (Myomaps, WIP1048 and WIP1041B respectively). Below is a more detailed description of the sequences used:

- IR-TSE ^22^: An inversion recovery turbo spin echo (IR-TSE) sequence was used as a reference to measure T1 values. Maps were generated offline using a custom program (Matlab, The MathWorks, Inc., Natick, MA, USA) based on six images collected using slice selective IR with TI=33, 100, 300, 900, 2700, 5000 ms; TE/TR=12ms/10s; slice thickness 8mm; flip angle = 90°; matrix 192×144; FOV~360×270mm and turbo factor=7. We performed a paired t-test to ensure that TF=7 did not give different T1 values from conventional IR-SE (TF=1).
- MC-SE: A multi contrast spin echo sequence was used to measure T2 values. Maps were generated offline using in-house software (Matlab, The MathWorks, Inc., Natick, MA, USA) based on 32 images collected using different echo times with TE/ =13.2/13.2/3000 ms; slice thickness 5mm; flip angle = 180°; matrix 256×184; FOV~360×270mm.
- MOLLI 5(3)3 ^10^: 8 SSFP images, TE/TR = 1.07/275 ms; minimum TI = 100ms; TI increment = 80 ms; slice thickness = 8mm; flip angle = 35°; matrix 192×144; FOV~360×270mm and GRAPPA acceleration factor = 2.
- ShMOLLI ^11^: 7 SSFP images; TE/TR = 1.07/275 ms; minimum TI = 100ms; TI increment = 80 ms;; slice thickness = 8mm; flip angle = 35°; matrix 192×144; FOV~360×270mm and GRAPPA acceleration factor = 2.
- SASHA ^12^: 11 SSFP images; TE/TR = 1.07/912 ms; minimum TI = 82 ms; TI increment = 79 ms; slice thickness = 8mm; flip angle = 70°; matrix 192×144; FOV~360×270mm and GRAPPA acceleration factor = 2.
- MTR ^23^: Magnetization transfer ratio maps were calculated using spoiled gradient echo acquisitions (FLASH) with (MT_on) and without (MT_off) off-resonance MT pulses. The FLASH sequence was used with TE/TR = 12/35 ms; flip angle = 5 deg; FOV ~360×270mm, acquisition matrix 320×256 and 8mm slice thickness. MTR was computed as MTR=(1 - MT_on/MT_off)* 100(%).

### Image analysis

For the ex vivo hearts, T1, T2 and MTR values were reported after manually delineating the endocardial and epicardial contours and generating masks of the LV myocardium on the parametric map using CVI42 (Circle CVI Inc., Calgary, Canada). All maps were resampled and aligned to match the MOLLI, ShMOLLI and SASHA outputs generated by the scanner. For each time point, we calculated MTR (1 – MT_on/MT_off), T2 (using CVI42), IR-TSE T1 map (code found here), MOLLI (Myomaps), ShMOLLI (WIP1048), SASHA (WIP1041B), and observed how the computed values changed over time. At every time point, the T1 bias was computed as the difference between MOLLI, ShMOLLI, SASHA and the reference IR-TSE T1 value.

### Statistical Analysis

Scatter plots were inspected to evaluate whether bivariate distributions of T1 bias vs MTR and T1 bias vs T2 effects display a marked nonlinearity and discernable outliers. Relationships appeared to follow a monotonic linear trend, although salient outliers were present for some of the plots. Moreover, some distributions exhibited heteroscedastic (fan-shaped) scatter. It is known that such distributions and outliers may pivot the linear regression line and misinterpret the strength of linear association between bivariate pairs. These effects can be mitigated using robust correlation analysis ^24^.

Based on the initial inspection of data, an open source toolbox by Pernet et al. (2013) ^24^ was employed on MATLAB R2015b (Mathworks Inc., Natick, CA) to calculate skipped correlations i) between pairs of T1 bias vs MTR and T1 bias vs T2 for MOLLI, ShMOLLI and SASHA and ii) between pairs of T1 vs MTR and T1 vs T2 for reference IR-TSE. A skipped correlation is calculated by first detecting and discarding the outliers from the best-fit distribution using the box-plot rule, where outliers are defined as those falling approximately three interquartile ranges (a measure of dispersion) above or below the bivariate distribution. Next, Pearson’s correlation and associated p-values were calculated on the remainder of the data by taking the sample size before outlier removal into account to control false positive rates. As the last step, 95% upper and lower confidence intervals (CIs) per skipped correlation were calculated using percentile bootstrapping, where inclusion of the zero in CIs indicates insignificant association between tested pairs. Statistical significances of skipped correlations in this study were determined based on this criterion (insignificant if the CI includes zero). Bootstrapped CI’s are used here because the traditional p-values are not protective against deviations from a distribution with uniform variance (homoscedasticity) and are more likely to produce false positive errors. Note that the computation of robust correlation effects requires a minimum sample size of 10, making skipped-correlation inapplicable to the phantom data. Therefore, for this dataset, the strength of relationship was determined using Pearson’s correlation without outlier detection. In the absence of this skipped-correlation test, significance for phantom data was evaluated by bootstrapped CIs to make interpretation less sensitive to the influence of data points deviating from the central envelope of the distribution. To determine whether the metrics changed significantly before and after fixation, we performed a separate Wilcoxon signed rank test on each of the metrics (T1, T2, MTR, T1 bias). Finally, differences between paired correlations before and after fixation were tested for significance by calculating CIs using percentile bootstrapping.

### Reproducibility

We believe that transparency is essential for the advancement of quantitative MRI, and that free dissemination of imaging data and analysis will help create a consensus in the field of qMRI, bringing us one step closer to clinical use. To that effect, we have made a strong effort to make all of the data, analysis and figures as transparent as possible. The data can be accessed via the <u>Open Science Framework</u>, and from there one can run the corresponding analysis and generate the figures included at the end of this manuscript. Additionally, our statistical framework has previously been published as an open-source <u>toolbox</u>, and the data visualization uses interactive figures (Plotly Technologies Inc., Montreal, Canada) to enable readers and reviewers to explore the data. The qMRI fitting code is currently not part of the documentation, in large part because the mapping routines are embedded in the scanner post-processing software and not publicly available.

## RESULTS

### Phantom

The correlations for the T1 bias against T2 and MTR in the agar phantom are shown in Figure 1, where each marker represents the average metric (T1 bias, T2, MTR) over the corresponding tube. The CIs for all correlations of the MOLLI_T1bias_ and ShMOLLI_T1bias_ did not include zero, indicating significant correlations with T2 and MTR (r < -0.85). On the other hand, the correlations between SASHA_T1bias_ and T2 and MTR were weak and insignificant, as indicated by quite wide CIs that include zero, and low correlation coefficients (r = 0.26). Overall, these findings point out a markedly high sensitivity of MOLLI_T1bias_ and ShMOLLI_T1bias_ to the T2 and MTR in comparison to that of SASHA_T1bias_ in phantoms.

**Figure 1.**
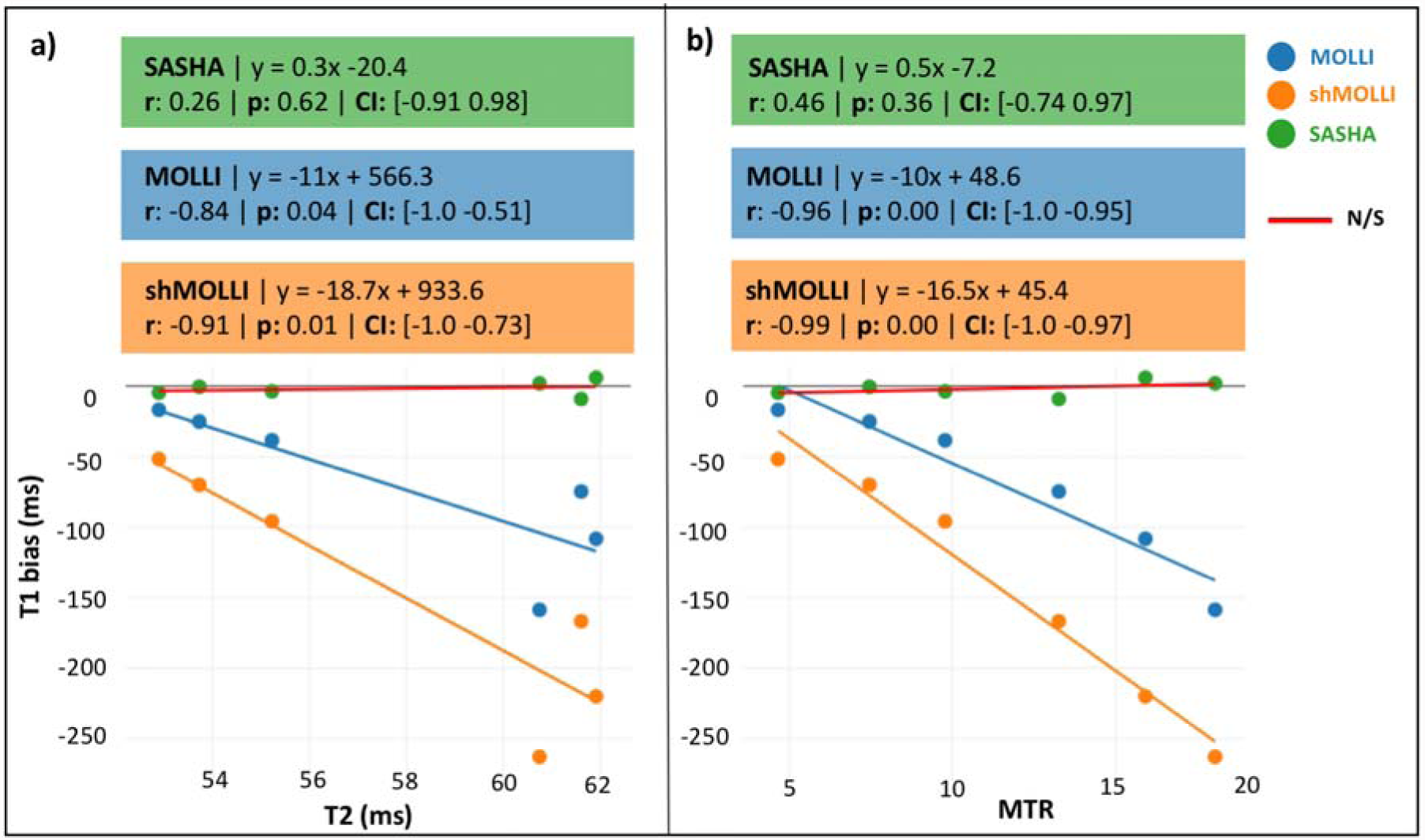
T1_bias_ correlations between cardiac T1 mapping sequences MOLLI (blue), shMOLLI (orange), SASHA (green) and **a)** T2 and **b)** MTR in the agar phantoms. Color-coded annotation boxes display the best-fit line equation, Pearson’s correlation coefficient (r) and its associated p-value, as well as the bootstrapped confidence intervals (CI). Red best fit lines indicate that the corresponding confidence interval includes zero, and is therefore not significant. Interactive version of this figure is available at http://neuropoly.pub/pigHeartsInteractive.

### Ex vivo

Upon 10% formalin fixation, a significant decrease was induced in the measured T1, T2 and MTR (Table 1). Although the SASHA_T1bias_ exhibited a significant decrease of 73% after fixation, changes in MOLLI_T1bias_ and ShMOLLI_T1bias_ were substantially smaller (Fig. 2b). Fixation also caused a notable increase in the variability of myocardial T1 distributions for all sequences. Figure 3 shows this for a representative specimen (heart #2), using box-and-whisker plots reporting the myocardium T1 values.

**Figure 2.**
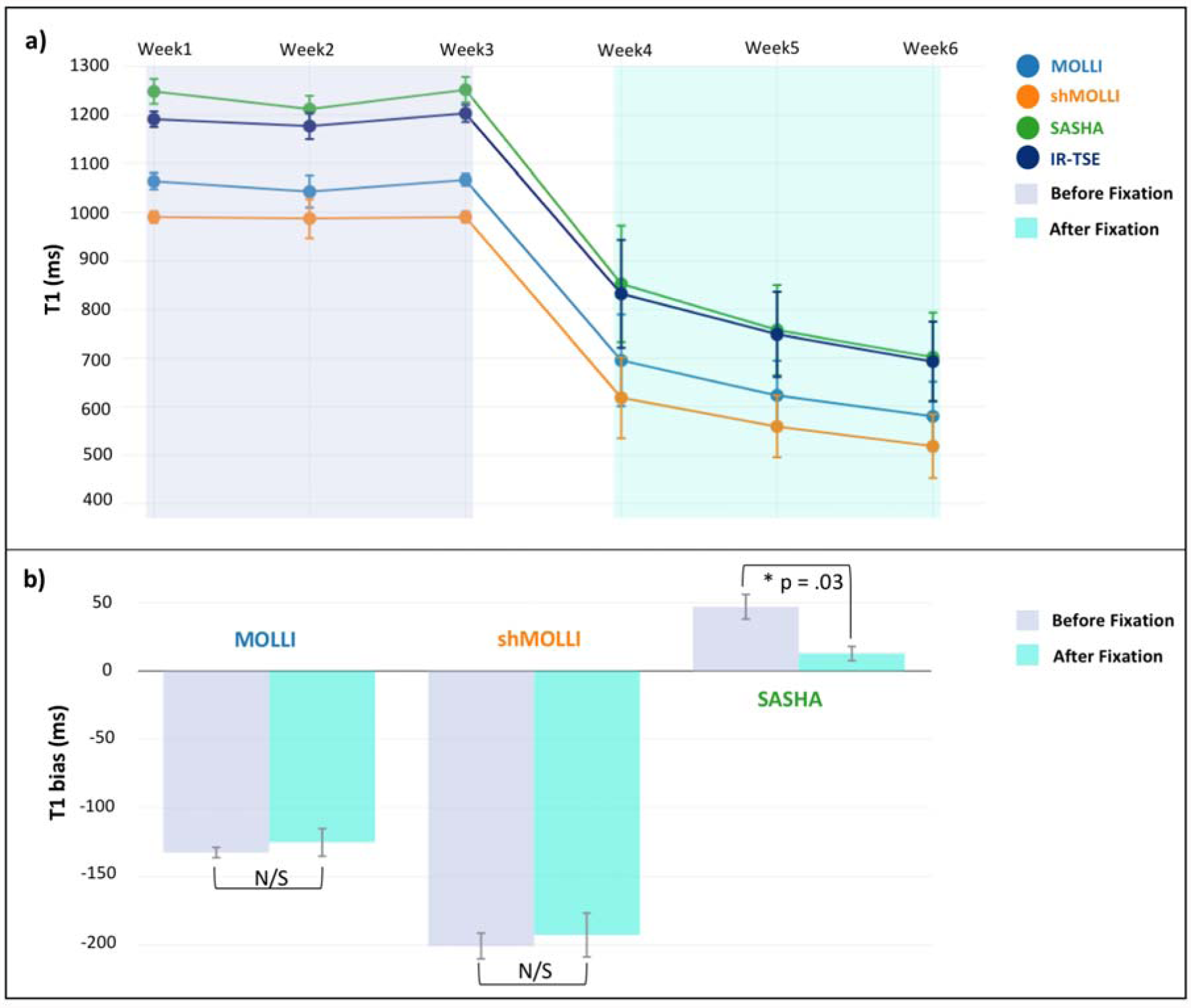
**a)** Evolution of T1 over time for all four sequences. MOLLI and SHMOLLI consistently underestimate, whereas SASHA overestimates the reference T1 values as estimated by IR-TSE. **b)** Comparison of T1 bias before (lavender) and after fixation (turquoise). Among the cardiac T1 mapping sequences, only the SASHA_T1bias_ shows a fixation-induced significant change. Statistical significance was evaluated by Wilcoxon signed rank test between pairs, after removing outlier effects. Interactive version of this figure is available at http://neuropoly.pub/pigHeartsInteractive.

**Figure 3.**
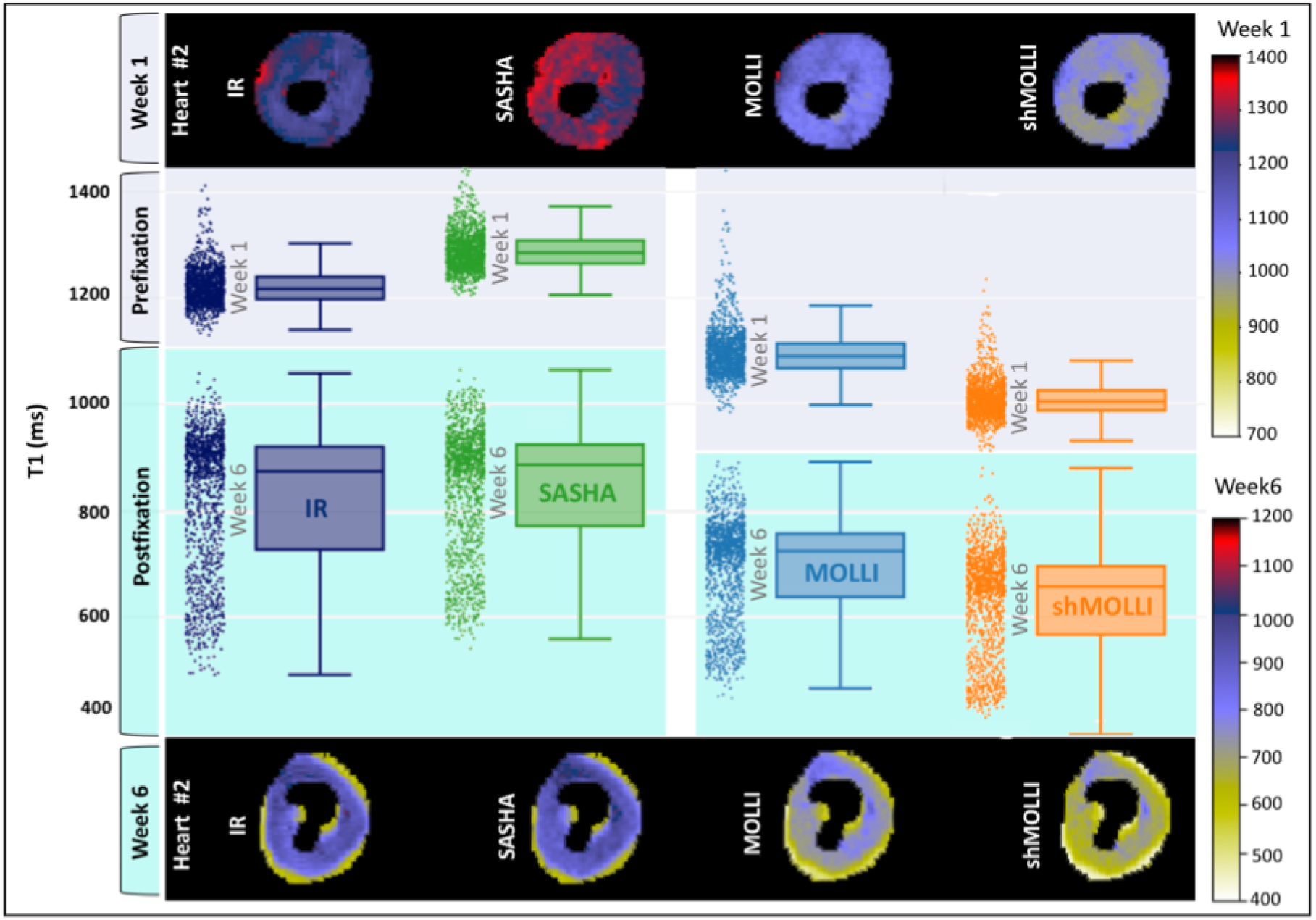
Voxelwise distributions of the myocardial T1 values (IR_T1_, SASHA_T1_, MOLLI_T1_, SHMOLLI_T1_) of a representative heart sample are shown by boxplots coupled with jittered markers in the middle panel for the first week (prefixation, lavender) and the sixth week (postfixation, turquoise). The upper and lower panels illustrate segmented myocardium T1 maps for the first week and the sixth week, respectively. Interactive version of this figure is available at http://neuropoly.pub/pigHeartsInteractive.

**Table 1:** The effect of 10% formalin fixation on the quantified metrics.

| **Metric** | **Prefixation**<br>**average** | **Postfixation**<br>**average** | **Change**<br>**%** | **p-value** |
| --- | --- | --- | --- | --- |
| *IR<sub>T1</sub> | 1190.4 ms | 758.09 ms | 36.31 % | < 0.001 |
| *MOLLI<sub>T1</sub> | 1057.90 ms | 632.86 ms | 40.18 % | < 0.001 |
| MOLLI<sub>T1bias</sub> | -132.44 ms | -125.23 ms | 5.44 % | 0.30 |
| *ShMOLLI<sub>T1</sub> | 989.45 ms | 565.30 ms | 42.86 % | < 0.001 |
| ShMOLLI<sub>T1bias</sub> | -200.93 ms | -192.79 ms | 4.05 % | 0.61 |
| *SASHA<sub>T1</sub> | 1237.4 ms | 770.78 ms | 37.70 % | < 0.001 |
| *SASHA<sub>T1bias</sub> | 46.986 ms | 12.69 ms | 72.99 % | < 0.001 |
| *MTR | 40.44 % | 33.66 % | 16.76 % | < 0.005 |
| *T2 | 71.949 ms | 59.58 ms | 17.18 % | < 0.001 |

Figure 4a-c shows that the reference T1 values before fixation do not show a significant correlation with T2, nor with MTR (r = -0.16 and r = 0.26, respectively). Even though fixation induced significant changes in T2, the strength of the relationship between IR_T1_ and T2 remained insignificant (See Fig. 4b and Table 2). Interestingly, while MTR was not correlated with the reference T1 before fixation (Table 1), formalin treatment gave rise to a significant correlation of 0.68 between IR_T1_ and MTR (Fig. 4d). The change between the paired correlations was also significant (Table 2).

**Figure 4:**
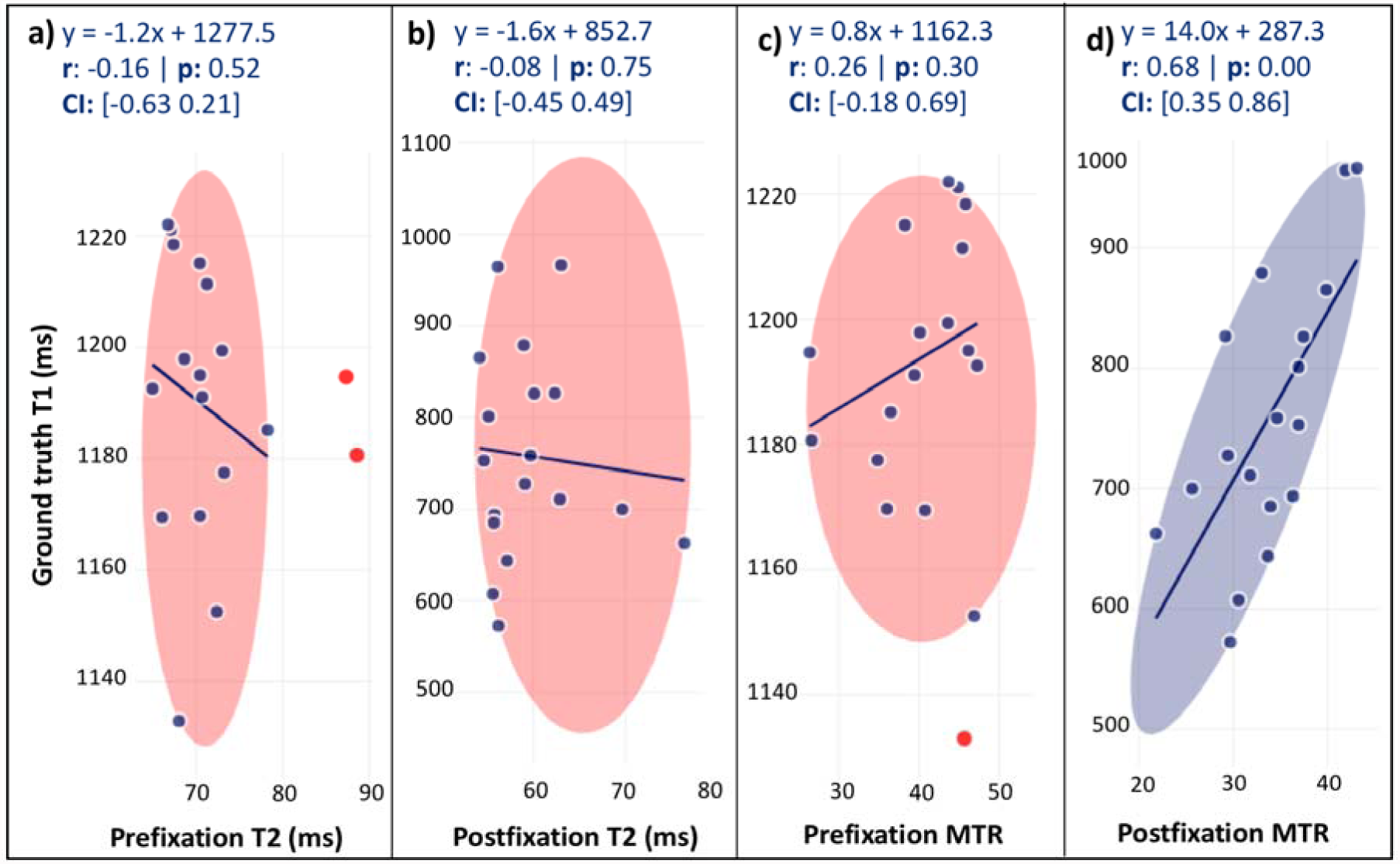
Correlation between IR_T1_ and T2 (a, b) and IR_T1_ and MTR (c, d) are calculated by skipped correlation before (a, c) and after fixation (b, d) in explanted pig hearts. Best fit line equation, Pearson’s correlation coefficient (r) and its associated p-value, as well as bootstrapped confidence intervals (CI) are displayed at the top of each panel. Red markers indicate bivariate outliers detected by the boxplot rule. Non-outlying points are encircled by an ellipse. The ellipse is red if the association determined by skipped correlation is insignificant (CI includes zero). Interactive version of this figure is available at http://neuropoly.pub/pigHeartsInteractive.

**Table 2.** Correlation of native myocardial T1 values (SASHA_T1_, MOLLI_T1,_ ShMOLLI_T1_) with T2 and MTR, before and after fixation. Significance of the differences between (before and after fixation) correlations were calculated using percentile bootstrapping.s

**Native T1 vs T2 Correlations**
|  | Prefixation |  |  | Postfixation |  |  | Difference |  |
| --- | --- | --- | --- | --- | --- | --- | --- | --- |
|  | r | p | 95% CI | r | p | 95% CI | r | 95% CI |
| **MOLLI** | -0.04 | 0.88 | [-0.47 0.55] | 0.11 | 0.66 | [-0.31 0.63] | -0.15 | [-0.74 0.69] |
| **ShMOLLI** | 0.16 | 0.53 | [-0.37 0.66] | -0.01 | 0.96 | [-0.37 0.53] | 0.17 | [-0.59 0.81] |
| **SASHA** | -0.35 | 0.16 | [-0.72 0.02] | -0.10 | 0.68 | [-0.5 0.52] | -0.24 | [-0.92 0.23] |
| **IR-TSE** | -0.16 | 0.52 | [-0.63 0.21] | -0.08 | 0.75 | [-0.45 0.49] | -0.08 | [-0.67 0.36] |

**Native T1 vs MTR Correlations**
|  | Prefixation |  |  | Postfixation |  |  | Difference |  |
| --- | --- | --- | --- | --- | --- | --- | --- | --- |
|  | r | p | 95% CI | r | p | 95% CI | r | 95% CI |
| **MOLLI** | 0.12 | 0.63 | [-0.30 0.54] | 0.67* | 0.00 | [0.32 0.85] | -0.55* | [-0.97 -0.06] |
| **ShMOLLI** | -0.49* | 0.04 | [-0.73 -0.13] | 0.62* | 0.01 | [0.18 0.84] | -1.11* | [-1.43 -0.62] |
| **SASHA** | 0.38 | 0.12 | [-0.02 0.86] | 0.69* | 0.00 | [0.36 0.87] | -0.32 | [-0.77 0.22] |
| **IR-TSE** | 0.26 | 0.30 | [-0.18 0.69] | 0.68* | 0.00 | [0.35 0.86] | -0.42* | [-1.00 -0.02] |

Consistent with the phantom data, the ex vivo robust correlation analysis showed that the correlation between SASHA_T1bias_ and T2 was weak and insignificant both before and after fixation (Fig. 5a-b). Unlike the SASHA_T1bias,_ the MOLLI_T1bias_ and ShMOLLI_T1bias_ exhibited much higher correlations with T2, also consistent with our phantom observations.

**Figure 5:**
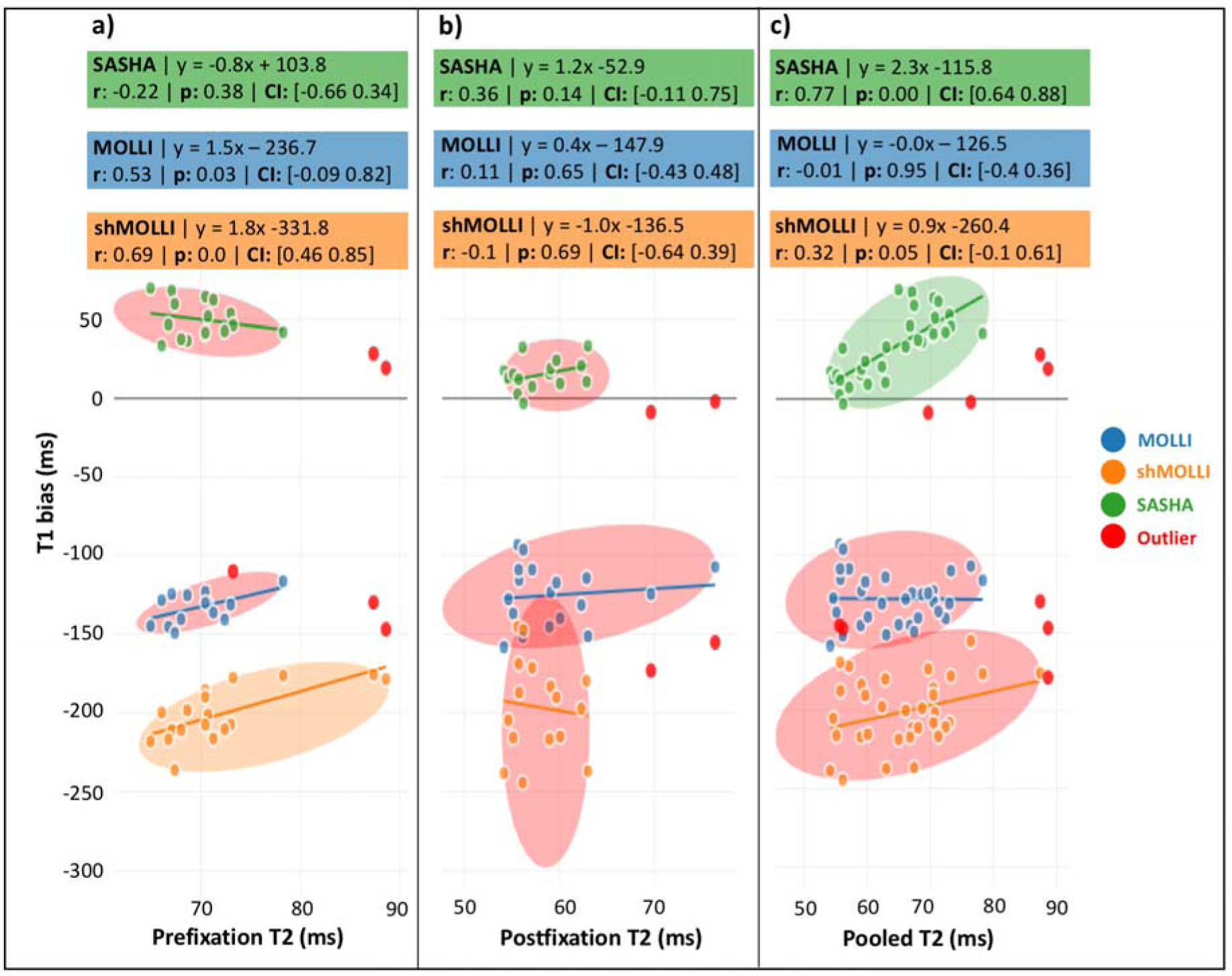
Correlation of T1 bias with T2 in ex-vivo pig myocardium is displayed for MOLLI, shMOLLI and SASHA **a)** before fixation, **b)** after fixation and **c)** pooled. Best fit line equation, Pearson’s correlation coefficient (r) and its associated p-value, as well as bootstrapped confidence intervals (CI) are displayed at the top of each panel. Red markers indicate bivariate outliers detected by the boxplot rule. Non-outlying points are encircled by an ellipse. The ellipse is red if the association determined by skipped correlation is insignificant (CI includes zero). Interactive version of this figure is available at http://neuropoly.pub/pigHeartsInteractive.

While T2 did not account for the bulk of the T1 bias, MTR was highly correlated with T1 bias for all cardiac T1 mapping sequences (Fig. 6). More importantly, the formalin treatment only slightly reduced the correlations between T1 bias and MTR (see Fig. 6a-b) and none of the differences between these paired correlations were significant.

**Figure 6:**
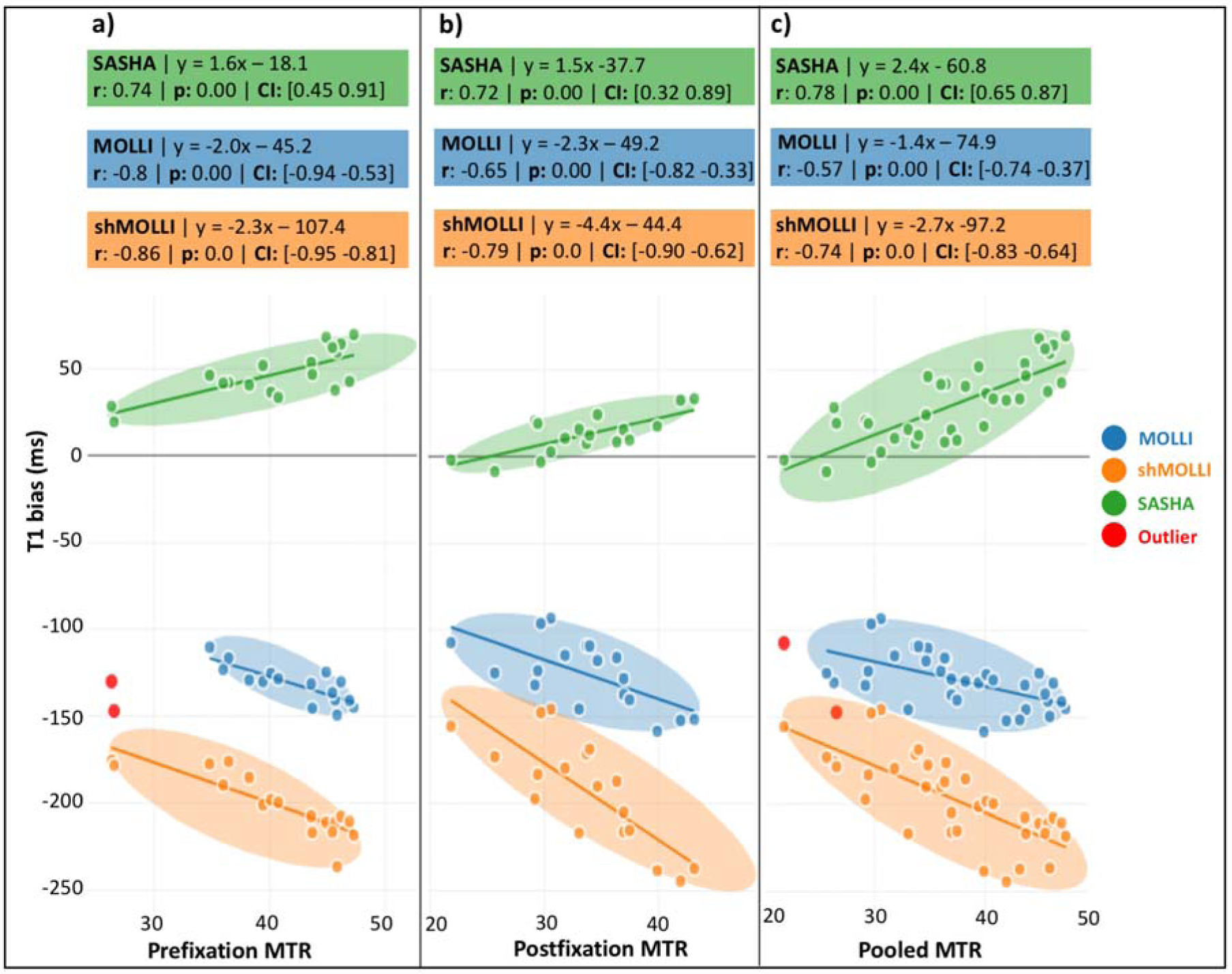
Correlation of T1 bias with MTR in ex-vivo pig myocardium is displayed for MOLLI, shMOLLI and SASHA **a)** before fixation, **b)** after fixation and **c)** pooled. Best fit line equation, Pearson’s correlation coefficient (r) and its associated p-value, as well as bootstrapped confidence intervals (CI) are displayed at top of each panel. Red markers indicate bivariate outliers detected by boxplot rule. Non-outlying points are encircled by an ellipse. The ellipse is red if the association determined by skipped correlation is insignificant (CI includes zero). Interactive version of this figure is available at http://neuropoly.pub/pigHeartsInteractive.

## DISCUSSION

We investigated cardiac T1 mapping sequences in a setting that allowed us to explore their accuracy and their dependence on T2 and magnetization transfer effects. The T2 effects were not significant (Fig. 5), and could not account for the T1 bias of MOLLI, ShMOLLI, SASHA with respect to the reference. On the other hand, the T1 biases exhibited a strong correlation with MTR (Fig. 6), and this correlation holds both before and after fixation. We conclude that inaccuracies in cardiac T1 mapping are primarily due to MT effects.

Differences between cardiac T1 mapping sequences have been reported before ^14^, and these differences are often higher in vivo than in phantoms ^14,25^. This is most likely due to the complex microstructure of human tissue, where MT effects are more complex. Formalin fixation gave us a sufficient dynamic range to explore realistic variations in the heart, but it also alters the biophysical properties of tissue ^26,27^, which is why we cannot conclude that the same observations will hold in vivo and under pathological conditions.

We saw that the inversion recovery sequences (MOLLI and ShMOLLI) exhibited higher correlations with MT and T2, whereas SASHA, despite being a saturation recovery sequence, followed the reference measurement more closely. On the other hand, the raw SASHA images have lower SNR, resulting in less uniform T1 maps, suggesting a trade-off between accuracy and precision. Both need to be taken into account before deciding on a specific T1 mapping protocol.

The correlation of T1 bias with MTR was prominent irrespective of the fixation state for all cardiac T1 mapping techniques. Following the significant decrease in MTR upon fixation (Table 1), correlation of IR_T1_ with MTR changed from a negligible to a significant level (Fig.4). However, a similar change in the correlation between T2 and IR_T1_ is not observed. Therefore, what remains after subtracting reference IR_T1_ from MOLLI, ShMOLLI and SASHA can be taken as a metric to evaluate the tradeoff between accuracy and MT sensitivity.

Given that relaxation processes and MT are naturally intertwined and subjected to complex alternations depending on the environment of water protons ^28,29^, phantoms and Bloch simulations could serve as the starting point for bringing the sequences together. Bloch simulations by Robson et al. reported at least a 10% reduction of the MOLLI_T1_ due to MT linked effects ^30^. They also showed that MOLLI is more sensitive to T2 and MT in comparison to SASHA. Our phantom and ex-vivo findings confirm this previously theoretically shown effect. Moreover, a recent study has drawn further attention to the MT effects in myocardial relaxometry quantification by combining MT informed simulations with phantom measurements^31^. In that study, Xanthis et al. tailored the MOLLI sequence to make it more T2, but less MT sensitive, which increased T1 estimation accuracy when compared to conventional MOLLI. This improvement provides support to our claim that MT contributes to T1 accuracy more than T2. While these effects are less pronounced in phantoms, it is critical to account for them in vivo in order to ensure proper comparison across sites, protocols and vendors.

## CONCLUSION

Our ex-vivo analysis brings us closer to a unified theory of cardiac T1 mapping, but additional histological, statistical and clinical analyses are necessary to create a T1 mapping consensus in the CMR community. We believe that transparency is essential for the advancement of quantitative MRI, and that free dissemination of imaging data and analysis will help create a consensus in the field, bringing us one step closer to routine use of qMRI in the clinic. This manuscript is our contribution to these efforts.

## ACKNOWLEDGEMENT

We thank the Montreal Heart Institute Foundation, Natural Sciences and Engineering Research Council (06774-2016), Fonds de recherche Santé Québec (36759 and 35250), the Quebec Bioimaging Network and the Canadian Open Neuroscience Platform for funding this study.

